# Antigen-specific T cell immunotherapy by in vivo mRNA delivery

**DOI:** 10.1101/2024.10.29.620946

**Authors:** Fang-Yi Su, Jamison C. Siebart, Ching S. Chan, Matthew Y. Wang, Xinyi Yao, Aaron Silva Trenkle, Avanti Sivakumar, Melanie Su, Rustin Harandi, Neha Shahrawat, Chi H. Nguyen, Anshika Goenka, Jinhee Mun, Madhav V. Dhodapkar, Gabriel A. Kwong

## Abstract

Immunotherapy has shown promise for treating patients with autoimmune diseases or cancer, yet treatment is associated with adverse effects associated with global activation or suppression of T cell immunity. Here, we developed antigen-presenting nanoparticles (APNs) to selectively engineer disease antigen (Ag)-specific T cells by *in vivo* mRNA delivery. APNs consist of a lipid nanoparticle core functionalized with peptide-major histocompatibility complexes (pMHCs), facilitating antigen-specific T cell transfection through cognate T cell receptor-mediated endocytosis. In mouse models of type 1 diabetes and multiple myeloma, APNs selectively deplete autoreactive T cells leading to durable control of glycemia, and engineer virus-specific T cells with anti-cancer chimeric antigen receptors (CARs), achieving comparable therapeutic outcome as virally transduced *ex vivo* CAR. Overall, our work supports the use of APNs to engineer disease-relevant T cells *in vivo* as Ag-specific immunotherapy for autoimmune disorders and cancer.

## Main

*In vivo* cell engineering is transforming immunotherapy by enabling direct manipulation of cellular fate and functions within the body^1, 2, 3, 4^. This approach bypasses the complicated *ex vivo* manufacturing process used for engineered cell therapies, such as T cells modified with anti-cancer chimeric antigen receptors (CARs)^5, 6^ and stem cell-derived pancreatic islet cells for type 1 diabetes (T1D)^7^. Approaches for *in vivo* cell engineering are built on advances in gene delivery vehicles, including virus-based vectors and non-viral nanoparticles^8^. For instance, viral vectors have been developed to reprogram T cells^5, 6, 9, 10^, B cells^11^, macrophages^12^, and dendritic cells^13^ *in vivo*. While promising, viral vectors are limited by the risk of insertional mutagenesis, pre-existing or treatment-induced antiviral immunity, and stringent regulatory hurdles^14, 15, 16^. Alternatively, mRNA delivered by lipid nanoparticles (LNPs) presents advantages in both safety and manufacturing, as mRNA translation does not require transgene insertion to the cell genome, and the safety profile and scalability of LNPs are evidenced by LNP-based COVID-19 vaccines^17^. The tropism of LNPs can be directed by surface modification of antibodies to preferentially deliver modulatory mRNA to selected cell populations, such as αCD117 to deliver pro-apoptotic mRNA to HSCs and pan-T cell antibodies (e.g., αCD3, αCD5) to deliver anti-cancer CAR mRNA to circulating T cells^3, 4, 18^. These examples highlight the growing interest in *in vivo* cell engineering to reap the full potential of engineered cell therapy.

While the estimated size of the human T cell repertoire exceeds 100 million, only a small subset of antigen (Ag)-specific cells is directly involved in the prevention or pathogenesis of a specific disease^19^. Therapeutic approaches that broadly target T cells (e.g., PD1 immune checkpoint and αCD3 antibody treatment) can result in global activation or suppression of T cell immunity, thereby making patients vulnerable to serious side effects, such as autoimmune diseases^20^, cytokine release syndrome^21^, and opportunistic infections^22^. *In vivo* engineering of disease-targeted, Ag-specific T cells, therefore, presents opportunities to enhance the specificity of T cell-based therapy. T cells recognize their cognate target cells through T cell receptors (TCRs), which interact with specific peptide antigens that are presented on major histocompatibility complex (MHC) molecules on the surface of the target cells. Emerging peptide-MHC (pMHC)-based technologies are being developed to modulate Ag-specific T cells *in vivo* with applications to enhance therapy for cancer and autoimmune diseases^23, 24, 25^. For example, Fc-fusion proteins and retrovirus modified with pMHCI molecules have been developed for delivery of immunomodulatory cytokines (e.g., IL2 and IL12) to expand and induce improved anticancer effects of cancer Ag-specific T cells^10, 26^. We recently developed Ag-presenting nanoparticles (APNs), which consist of lipid nanoparticles (LNPs) similar to the COVID-19 vaccines, but modified with pMHC molecules on the LNP surface through post-insertion^27^. APNs simultaneously delivered functional mRNA to multiple Ag-specific T cell subsets in TCR transgenic mouse models and a mouse model of human influenza infection.

Here we develop APN therapeutics for T1D, an autoimmune disease caused by autoreactive T cell-mediated destruction of pancreatic β cells^28^, and multiple myeloma (MM), a cancer of plasma cells in the bone marrow. We design APNs against two Ag-specific T cell populations by targeting autoreactive T cells to prevent the onset of hyperglycemia associated with T1D, and by engineering human antiviral T cells to produce CAR T cells for cancer treatment. In an adoptive transfer murine model of T1D, APNs that selectively deliver pro-apoptotic caspase 6 (Casp6) mRNA to autoreactive β-islet-specific CD8 T cells eliminate these cells to prevent the onset of hyperglycemia in a murine model of T1D while minimizing off-target T cell effects including liver and kidney toxicity. We further demonstrate *in vivo* re-programming of virus-specific T cells to express CAR mRNA to improve cancer treatment due to the memory phenotype of virus-specific T cells for prolonged *in vivo* persistence^29^ and stimulation of CAR T cells through their endogenous TCRs. Overall, our data support APNs to engineer disease-relevant T cells *in vivo* as Ag-specific immunotherapy.

## Results

### APN transfection of T cells is TCR-dependent

We previously developed a post-insertion method to formulate APNs by coincubating LNPs with pMHC molecules for *in vivo* transfection of Ag-specific T cells with reporter mRNA^30^. To further optimize the LNP core for mouse and human CD8 T cell transfection, we tested seven clinically approved or published formulations for transfecting primary mouse and human T cells with mRNA encoding a Nano-luciferase (nLuc) reporter (**Supplementary Table. 1**). While all seven LNP formulations transfected mouse and human CD8 T cells *in vitro* (**Supplementary Fig. S1**), we found that LNPs formulated with the ionizable lipid ALC-0315 showed the highest bioluminescence intensity in both mouse and human T cells (10^8^-10^9^ as compared to ∼10^6^ in the untransfected control). All APNs were therefore formulated with ALC-0315 for following studies.

Proteins involved in T cell reprogramming are typically secreted within the extracellular space (e.g., cytokines), expressed on the cell membrane (e.g., CAR), or localized within intracellular compartments (e.g., proapoptotic proteins). To test APN transfection of T cells with mRNA encoding for proteins within these compartments, we loaded APNs with mRNA encoding one of three reporters—secreted nLuc, fluorescent protein mTagBFP, or membrane-bound VHH nanobody (**Fig. 1A**). We functionalized APNs with K^d^/NRP-V7 pMHC molecules to transfect autoreactive Ag-specific T cells from transgenic NOD8.3 mice whose CD8+ T cells express a TCR that specifically recognizes NRP-V7 (KYNKANVFL), a peptide mimotope derived from the T1D-associated antigen IGRP^31^. We found that K^d^/NRP-V7 APNs transfected NOD8.3 CD8 T cells with all three mRNA constructs, whereas non-cognate APN treated groups showed minimal transfection (**Fig. 1B-D**). These data collectively support the activity of APNs in transfecting CD8 T cells with mRNA encoding for functional proteins localized to unique cellular compartments in an antigen-specific manner.

**Fig. 1.**
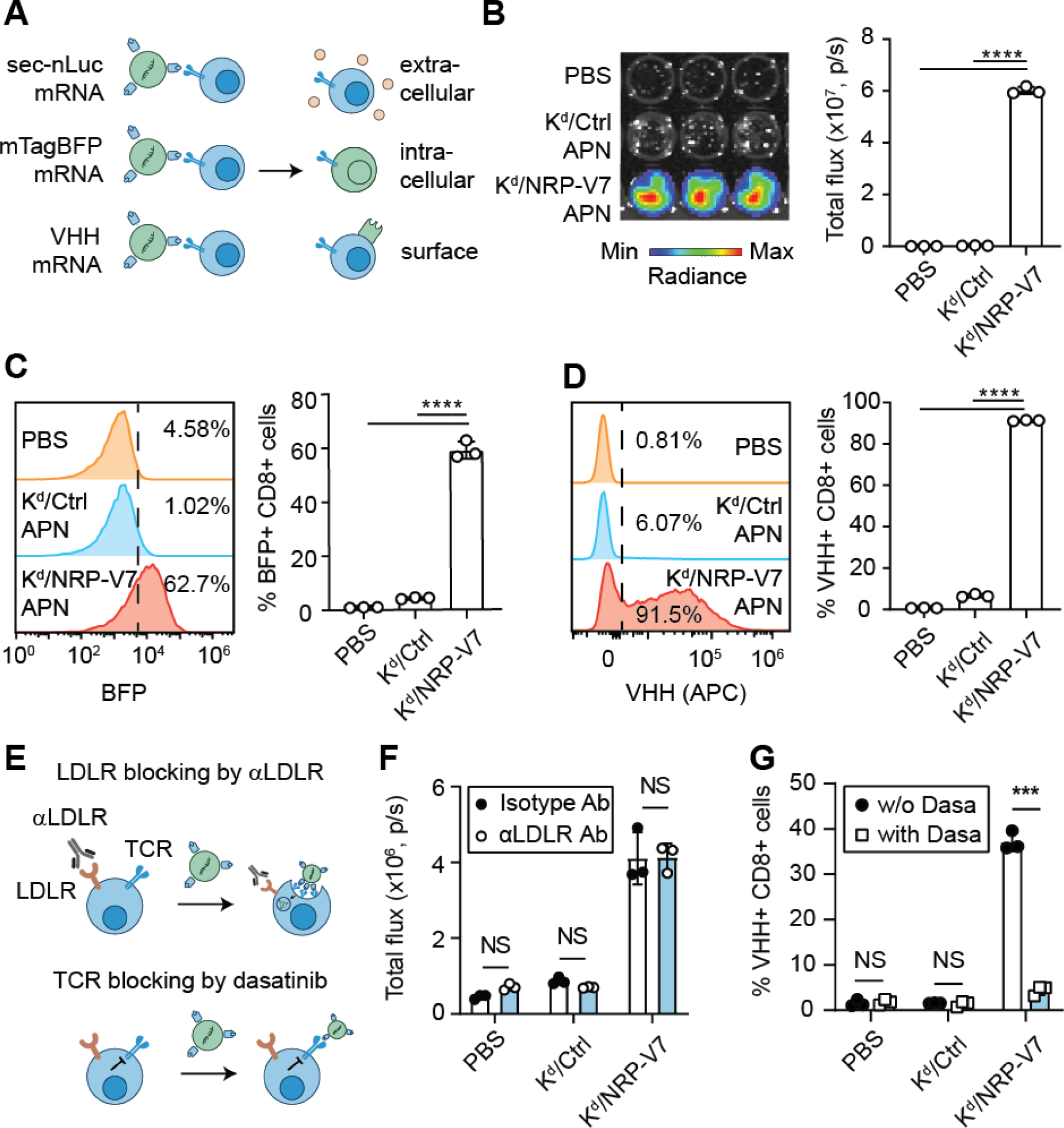
APNs transfect antigen-specific T cells in TCR-dependent manner. **(A)** Schematic showing APN delivery of diverse mRNA cargo to cognate T cells. b-g, Activated NOD8.3 CD8 T cells were transfected in vitro with PBS, K^d^/Ctrl (non-cognate control) APNs, or K^d^/NRP-V7 (cognate) APNs. After 24 hours, transfection readout was measured. **(B)** T cells transfected by APNs carrying secreted nLuc mRNA were analyzed via IVIS and quantified. **(C)** Representative flow plots and frequency bar plot of intracellular BFP expression. **(D)** Representative flow plots and frequency bar plot of surface-bound VHH expression. One-way analysis of variance (ANOVA) and Tukey post-test and correction for multiple comparisons; ****P<0.0001. All data are means ± SD; n=3 independent wells. **(E)** Schematic showing the internalization mechanism of APN by T cells through T cell receptor (TCR), not low-density lipoprotein receptor (LDLR). **(F)** Activated NOD8.3 CD8 T cells were coincubated with either a LDLR blocking antibody (αLDLR Ab) or an isotype antibody control (isotype Ab). APNs were encapsulated with nLuc and transfection was measured by IVIS 24 hours post transfection. **(G)** Quantification of APN transfection in the presence of the TCR signaling inhibitor dasatinib (dasa), measured via flow cytometry. Two-way ANOVA with Sidak post-test and correction for multiple comparisons. NS= not significant; ****P<0.0001.

We proceeded to understand the primary uptake mechanism of APNs by antigen-specific T cells. While LNPs are internalized by cellular endocytosis following association with ApoE and binding to the low-density lipoprotein receptor (LDLR)^32, 33^, pMHC tetramers induce TCR clustering and promote TCR-mediated endocytosis into T cells^34, 35^. We therefore, hypothesized that multivalent presentation of pMHC molecules on the APN surface could facilitate mRNA transfection through TCR clustering rather than through LDLR endocytosis (**Fig. 1E**). We found that pre-treatment of activated T cells with a LDLR blocking antibody (αLDLR) significantly reduced T cell transfection from bare LNPs compared to T cells treated with the isotype control antibody (**Supplementary Fig. S2**). In contrast to LNP transfection, APN-transfected T cells pretreated with αLDLR antibody or isotype control showed comparable luminescence signals (**Fig. 1F**), providing support that LDLR uptake was not the primary mechanism of APN transfection. We next treated T cells with dasatinib—a protein kinase inhibitor known to inhibit TCR signaling and prevent the internalization of TCR and bound pMHC multimers^35^—before and during APN transfection. In the presence of dasatinib, APN-transfected T cells exhibited reduced transfection efficiency compared to the untreated control (40% vs. 5%) (**Fig. 1G**). These results supported that APN-mediated transfection of T cells was dependent on TCR engagement rather than LDLR.

### APN selectively deplete cognate autoreactive T cells and maintain T cell homeostasis

T1D is an autoimmune disease in which autoreactive cytotoxic T cells destroy insulin-producing β cells within the pancreatic islets of Langerhans^36^. The FDA-approved αCD3 monoclonal antibody (teplizumab) delays the onset of clinical T1D by broadly suppressing the activity of CD8 T cells, including autoreactive CD8+ T cells^37, 38^. We therefore tested whether APNs could prevent the onset of T1D in an antigen-specific manner. To do this, we designed mRNA constructs encoding pro-apoptotic peptides (BIM, BID)^39^ or proteins (granzyme B, Casp9, and Casp6)^40, 41^ (**Supplemental Fig. 3A,B**) to prevent the onset of T1D by depleting autoreactive T cells. Using electroporation to deliver these mRNA constructs to primary mouse T cells, we screened and compared the efficiency of these pro-apoptotic mRNA constructs in inducing cell death (**Supplementary Fig. 3C**). We found that mouse Casp6 mRNA led to the most potent induced cell death and could be delivered through ALC-0315 LNPs to induce T cell death *in vitro* (**Supplementary Fig. 3D**). We therefore proceeded with this Casp6 for APN delivery in the following *in vivo* studies.

To test the ability of APNs to deplete therapeutically relevant T cells, we used a T1D mouse model by adoptively transferring peptide-pulsed NOD8.3 T cells into host wildtype NOD mice to induce hyperglycemia. We treated these mice with cognate APNs encapsulated with Casp6 (K^d^/NRP-V7 Casp6 APNs) to deplete these autoreactive T cells (**Fig. 2A**). We found that cell death was mRNA- and pMHC-dependent, as cognate APNs loaded with non-therapeutic mRNA (K^d^/NRP-V7 VHH APN) and non-cognate APNs loaded with Casp6 (K^d^/Ctrl Casp6 APN) depleted less NOD8.3 T cells in the blood, spleen, and pancreatic lymph nodes (pLN) compared to K^d^/NRP-V7 Casp6 APNs (**Fig. 2B,C**). As a positive control, we also compared APN treatment to Fc-nonbinding αCD3 (αCD3) treatment by following preclinical dosing schemes of 2.5 mg/kg doses daily for five days in a row^42^. We found that αCD3 treatment depleted NOD8.3 T cells (**Fig. 2B, C**), but was not antigen-specific, leading to decreased total CD8 T cell percentages in the blood, spleen, and pLN, while K^d^/NRP-V7 Casp6 APNs preserved the total CD8 T cell percentages (**Fig. 2D**). These results were in line with the initial lymphopenia observed in patients after αCD3 treatment^43^. Collectively, these data showed that APNs can selectively deliver Casp6 to deplete autoreactive T cells, while avoiding the reduction of total CD8 T cell percentages in the major organs and tissues.

**Fig. 2.**
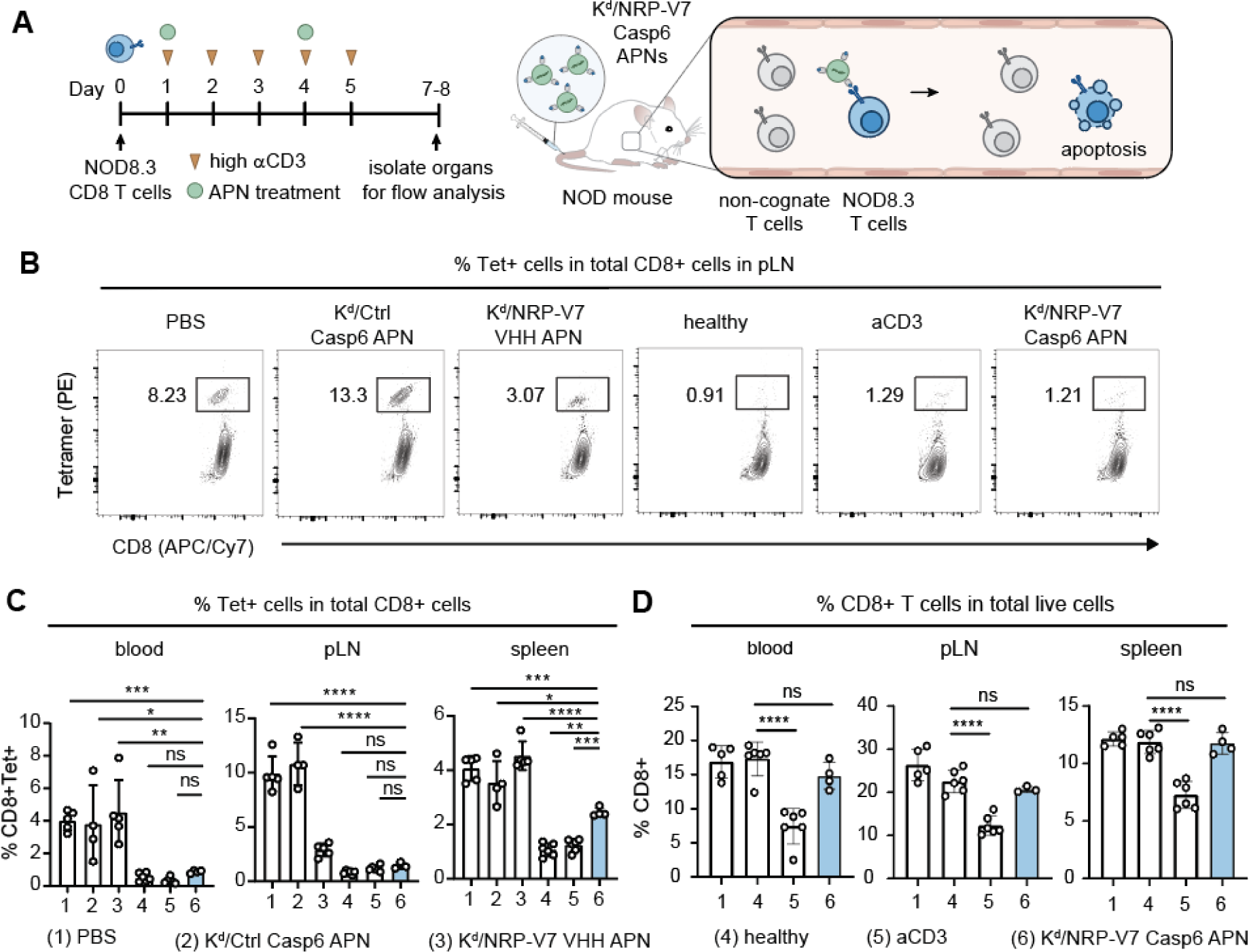
Casp6 APNs deplete autoreactive T cells in ACT model of T1D and maintain total T cell homeostasis. **(A)** Timeline describing T1D model development and treatment; APNs selectively target NOD8.3 T cells T cells in vivo and delivery of Casp6 mRNA triggers apoptosis. **(B)** Representative flow plots showing NOD8.3 T cells present in the pancreatic lymph node (pLN) after respective treatments. **(C)** Quantification of % NOD8.3 T cells in peripheral blood, pLN, and spleen after treatment. **(D)** Total CD8 T cells in the peripheral blood, pLN, and spleen after treatment. Organs isolated with less than 1% viable cells after processing were excluded from analysis. One-way ANOVA with Tukey’s post-test and correction for multiple comparisons, n=4-6 biological replicates. ns = not significant; *, **P < 0.01, ***P < 0.001, ****p<0.0001.

### APNs prevent the onset of hyperglycemia in a mouse model of type 1 diabetes

To assess the therapeutic relevance of APN-mediated depletion of autoreactive T cells, we tracked the blood glucose levels of host NOD mice after adoptive cell transfer of peptide-pulsed NOD8.3 T cells and treatment with APNs (**Fig. 3A**). We showed that mice treated with K^d^/NRP-V7 VHH APNs and K^d^/Ctrl Casp6 APNs had comparable high blood glucose levels as untreated mice (PBS) (> 250 mg/dL is considered diabetic) (**Fig. 3B,C**). However, mice treated with K^d^/NRP-V7 Casp6 APNs maintained healthy blood glucose levels like the αCD3 treated mice and healthy mice (<250 mg/dL), furthering demonstrating the necessity for both cognate pMHC for T cell uptake and relevant mRNA for translation of Casp6 protein to induce cell death (**Fig. 3B,C**). We also found that the prevention of T1D from K^d^/NRP-V7 Casp6 APNs was durable, as blood glucose were maintained at normal and stable levels up to 30 days after adoptive transfer of NOD8.3 T cells (**Fig. 3D**).

**Fig. 3.**
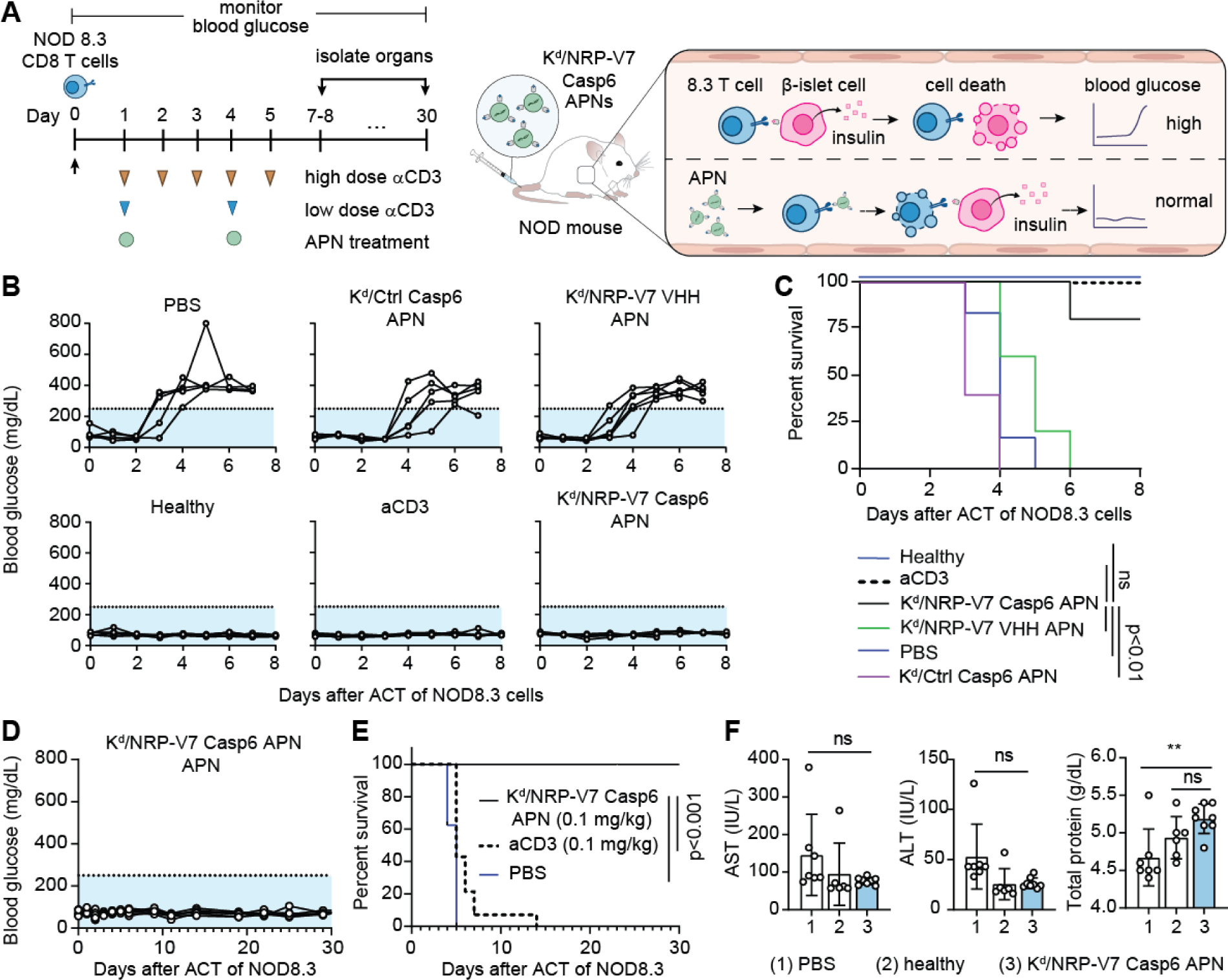
Casp6 APNs durably prevent onset of hyperglycemia and are well tolerated at given dose. **(A)** Timeline and schematic showing treatment strategies; low dose aCD3 and APN treatment was administered at 0.1 mg/kg and high dose αCD3 was administered at 2.5 mg/kg at the specified timepoints. APNs prevent hyperglycemia by selectively depleting autoreactive NOD8.3 T cells and sparing β-islet cell function. **(B)** Blood glucose traces of mice after receiving respective treatments; light-blue shaded region underneath dashed line represents healthy blood glucose levels (<250 mg/dL). **(C)** survival curve of mice; survival refers to living mice with blood glucose levels <250 mg/dL. Log-rank (Mantel-Cox) test, n=4-6 biological replicates, NS = not significant. **(D)** Long term blood glucose traces of Kd/NRP-V7 Casp6 APNs treated mice. **(E)** Survival curve of mice; survival refers to living mice with blood glucose levels <250 mg/dL. Log-rank (Mantel-Cox) test, n=8-16 biological replicates. **(F)** measure of liver enzyme alanine transaminase (ALT) and aspartate aminotransferase (ALT) levels and total protein levels in serum following administration of Kd/NRP-V7 Casp6 APNs. One-way ANOVA with Tukey’s post-test and correction for multiple comparisons; means ± SD, n=8-16 biological replicates. ns = not significant, **P<0.01.

We next sought to comparable the treatment efficacy of APNs to αCD3 antibody using comparable doses. When dosing schemes were matched head-to-head (i.e., 0.1 mg/kg dosed 1 day and 4 days after injection of NOD8.3 T cells), αCD3 treatment was only able to moderately delay the onset of hyperglycemia in this aggressive model (all mice became diabetic by day 14), while K^d^/NRP-V7 Casp6 APNs durably prevented the onset of hyperglycemia for at least 30 days (**Fig. 3E**). Additionally, K^d^/NRP-V7 Casp6 APN treatment was well tolerated at the given dose (0.1 mg/kg mRNA), with no observable changes in liver and kidney biochemical analyses (BUM, creatine, phosphorous, calcium) from blood serum compared to healthy mice (**Fig. 3F, Supplementary Fig. S4A**), and no observed decline in body weight of treated mice (**Supplementary Fig. S4B**). Taken together, these data demonstrated the potent activity of K^d^/NRP-V7 Casp6 APNs in preventing the onset of hyperglycemia without causing off-target toxicity in liver and kidney.

### APN transfect virus-specific T cells from MM patients with CAR *in vitro*

To test the anti-cancer potential of APN using a clinically relevant CAR construct and mouse models, we designed and validated a mRNA sequence encoding an αBCMA CAR construct similar to the ones tested in clinical trials (**Supplementary Fig. S5**)^44, 45^. We developed APNs to deliver αBCMA mRNA to HLA-A2.1+ human influenza A virus (IAV)-specific T cells with TCRs to recognize an immunodominant IAV peptide epitope (GILGFVFTL). Re-directing IAV-specific T cells with CAR leverages their memory phenotypes for prolonged *in vivo* persistence^29^ and enables stimulation of CAR T cells through their endogenous TCR using existing influenza vaccines to improve anti-tumor efficacy^46, 47, 48^. To do this, we incubated HLA/IAV APNs loaded with mRNA encoding either nLuc or αBCMA CAR with enriched human IAV-specific T cells *in vitro* for 24 hours. Compared to the untreated group, HLA/IAV APNs transfected IAV-specific T cells with nLuc to generate bioluminescence (∼100x higher in bioluminescence intensity compared to the untreated group) (**Fig. 4A**) and induced a dose-dependent CAR expression in IAV-specific T cells (50% and 65% at 1 and 2.5 μg mRNA doses per 10^6^ cells, respectively) (**Fig. 4B,C**). Notably, non-cognate T cells incubated with HLA/IAV APNs only showed αBCMA CAR expression comparable close to background levels. We next tested the effector functions of the APN-transfected αBCMA CAR T cells by co-incubating the CAR T cells with BCMA+ MM1R MM cancer cells that constitutively express renilla luciferase for evaluating cytotoxicity (**Fig. 4D**). After 24-hour co-incubation, T cells transfected by nLuc mRNA-loaded APNs resulted in comparable MM1R viability as the PBS control (∼100% viability) (**Fig. 4E**). By contrast, T cells transfected αBCMA CAR mRNA-loaded APNs resulted in significantly lower viability of MM1R (∼20%). These data indicate that APN can transfect human IAV-specific T cells with functional αBCMA CAR *in vitro*.

**Fig. 4.**
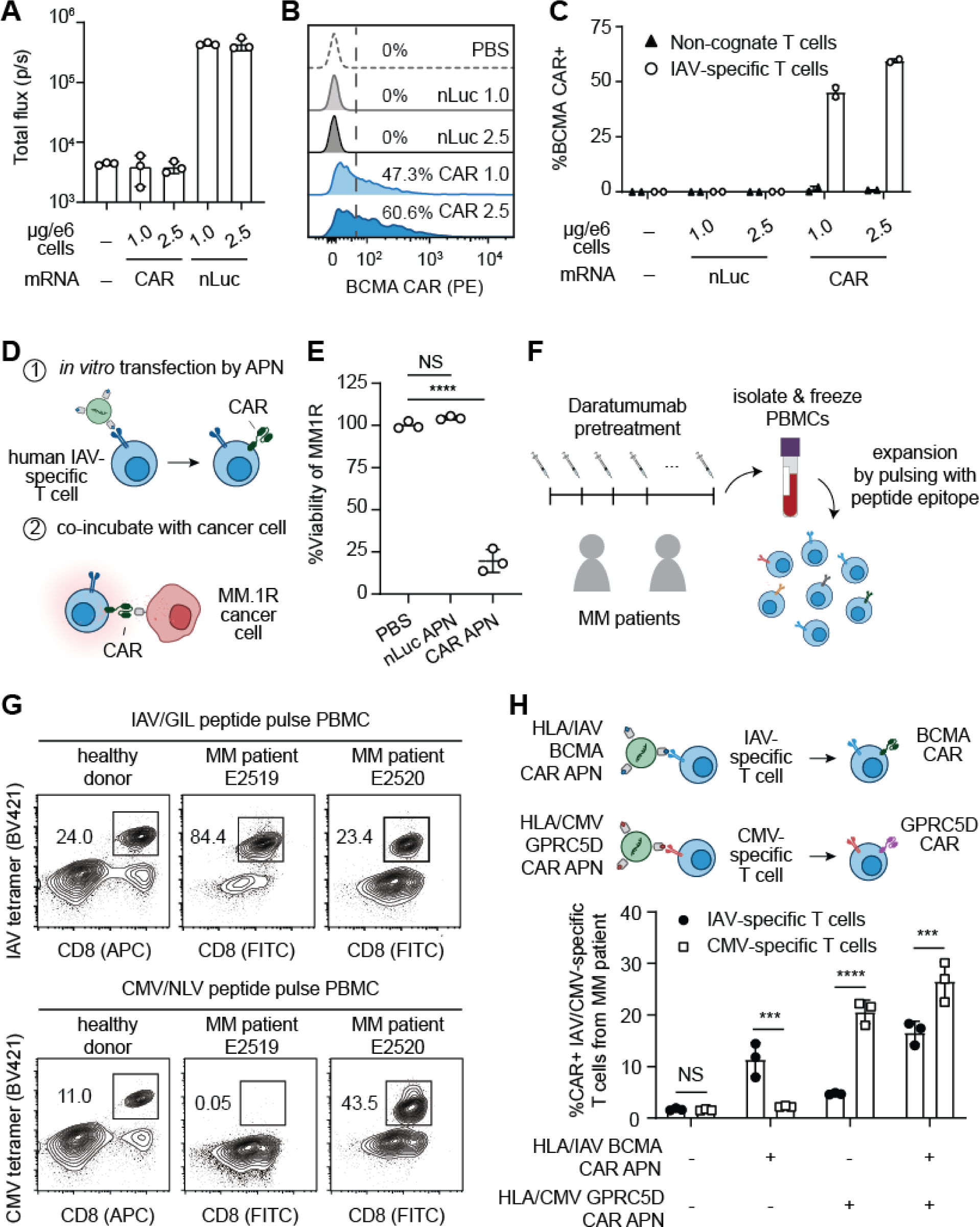
APN transfect human virus-specific T cells from a pretreated multiple myeloma patient with CAR mRNA *in vitro*. **(A)**-**(C)** Enriched human influenza A virus (IAV)-specific T cells from healthy donor were treated with APNs functionalized with human leukocyte antigen A2.1+ (HLA-A2.1) bound to IAV peptide (HLA/IAV APN). APNs were encapsulated with either nLuc or αBCMA CAR mRNA and transfection was analyzed through IVIS readout of nLuc **(A)** or flow cytometry detection of αBCMA CAR **(B)** to show transfection was limited to on-target IAV-specific T cells **(C)**. **(D)**-**(E)**, HLA/IAV APNs transfect human-IAV specific T cells with functional αBCMA CAR and kill luciferized MM.1R cancer cells in *in vitro* cocultures. **(D)** and quantified via bioluminescence IVIS readout **(E)**. One-way ANOVA with Tukey’s post-test and correction for multiple comparisons; means ± SD, n=3 independent wells. NS= not significant, ****P<0.0001. **(F)**-**(G)**, Frozen peripheral blood mononuclear cells (PBMC) were obtained from HLA-A2.1 positive MM patients who were previously treated with daratumumab and peptide pulsed with peptide epitopes **(F)**, leading to IAV and CMV-specific T cell expansion **(G)**. **(H)** Human MM patient CMV- and IAV-specific T cells were mixed together and transfected in vitro with HLA/IAV and HLA/CMV APNs encapsulated with αBCMA CAR and αGPRC5D CAR mRNA respectively. Two-way ANOVA with Sidak post-test and correction for multiple comparisons; means ± SD, n=3 independent wells. NS = not significant, ***P<0.001, ****P<0.0001.

The FDA-approved αBCMA CAR T cell therapies (Abecma, Carvykti) require patients to have undergone at least one prior line of therapy. Therefore, we sought to confirm that virus-specific T cells were not eliminated and could be expanded from patients with active MM, including those relapsing after initial therapy. Frozen PBMCs from two HLA-A2.1+ MM patients (E2519 and E2520) previously treated with daratumumab and a matching HLA-A2.1+ healthy donor (positive control) were pulsed with the immunodominant IAV peptide and cytomegalovirus (CMV) peptide NLVPMVATV to expand IAV-specific CD8 T cells and CMV-specific T cells, respectively (**Fig. 4F**). After 14 days, we detected robust expansion of IAV-specific T cells in both MM patients by tetramer analysis (85% and 23% positive compared to 24% from the healthy donor). We also observed robust expansion of CMV-specific T cells in MM patient E2520 (43.5% positive compared to 11% from the healthy donor), but not in MM patient E2519 (0.05%). These results collectively support IAV-specific and/or CMV-specific T cells are in circulation in pre-treated MM patients (**Fig. 4G**).

Leveraging the ability of APNs to simultaneously deliver mRNA to different antigen-specific T cell subsets *in vivo*^49^, we asked whether APNs could deliver two different CAR mRNA constructs to IAV-specific T cells and CMV-specific T cells, respectively (**Fig. 4H**). In addition to αBCMA CAR, we included a CAR construct targeting human GPRC5D (G-protein coupled receptor, class C, group 5, member D), which is also a highly expressed antigen by human MM cells^50, 51, 52^ and αGPRC5D CAR have shown promising therapeutic efficacy in clinical trials^53, 54^. Building on the αBCMA CAR mRNA construct we tested in **Supplementary Fig. S5**, we designed and validated a mRNA sequence encoding an αGPRC5D CAR construct^55, 56^ by replacing the scFv region (**Supplementary Fig. S6**). *In vitro,* we mixed an equal number of IAV-specific T cells and CMV-specific T cells and showed that mono-treatment with HLA/IAV αBCMA CAR APN preferentially transfected IAV-specific T cells, while HLA/CMV αGPRC5D APN selectively transfected CMV-specific T cells. Moreover, the combination of both HLA/IAV αBCMA CAR APN and HLA/CMV αGPRC5D APN resulted in CAR expression in both IAV-specific T cells and CMV-specific T cells (∼15–25% CAR+ respectively) (**Fig. 4H**). Those results indicate that APNs can transfect two virus-specific T cells isolated from a pretreated MM patient with CAR mRNA.

### APN-transfected CAR T cells reduce tumor burden *in* xenograft MM mouse model

We next assessed whether APNs loaded with BCMA CAR mRNA could reprogram IAV-specific T cells *in vivo*. We expanded immunodominant IAV-specific T cells by peptide pulse (GILGFVFTL) to mimic the IAV-specific T cell expansion after influenza vaccine (**Fig. 5A**). We infused immunodeficient NSG mice with 12 million IAV-peptide pulsed PBMC (containing ∼2 million IAV-specific CD8 T cells), resulting in IAV-specific CD8 T cells to be ∼1% of total splenocytes at 48 hours after the cell transfer to emulate the IAV-specific T cell frequency after vaccination in humans (∼0.5-1%) (**Fig. 5B**). At such a frequency, intravenous injection of HLA/IAV APNs to these NSG mice resulted in a dose-dependent CAR transfection efficiency (**Supplementary Fig. S7**) and achieved ∼40% transfection of IAV-specific CD8 T cells with αBCMA CAR at 1 mg/kg mRNA dose (**Fig. 5C**). By contrast, we observed no αBCMA CAR expression in NSG mice treated with non-cognate HLA/CMV APNs displaying a CMV peptide epitope. Similarly, PBS and APNs carrying αCD19 CAR mRNA did not result in detectable αBCMA CAR transfection in IAV-specific CD8 T cells.

**Fig. 5.**
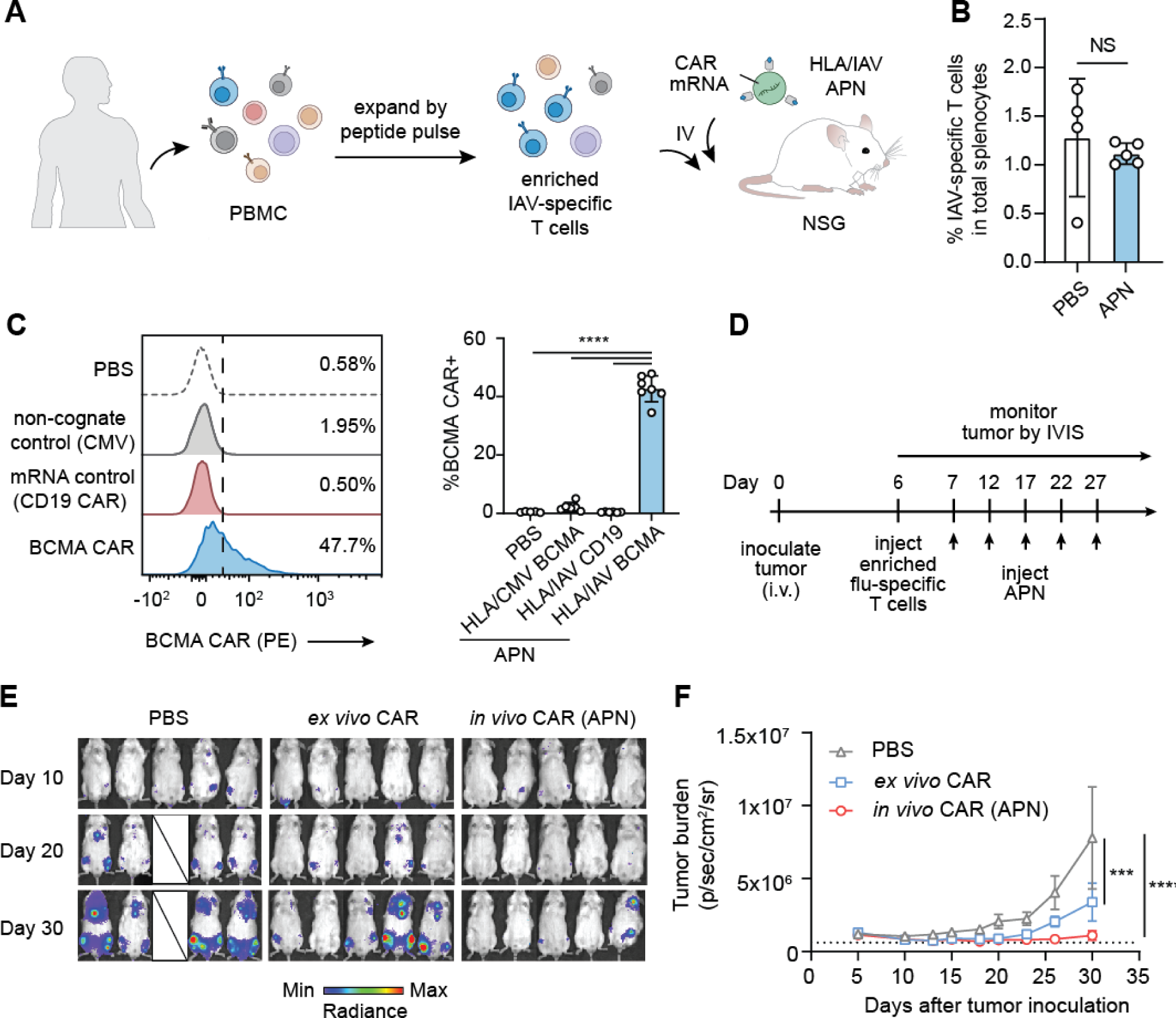
*In vivo* transfected αBCMA CAR T cells induce anti-cancer efficacy in NSG mice bearing systemic human multiple myeloma (MM). **(A)-(C)** Enriched human I nfluenza A virus (IAV)-specific T cells from a healthy donor were intravenously injected into NSG mice and dosed with HLA/IAV APNs carrying αBCMA CAR mRNA **(A)** and IAV-specific T cells in the spleen were quantified **(B)**. Student’s t-test; means ± SD, n=4-5 biological replicates. NS= not significant. (C) I*n vivo* transfection efficiency with HLA/IAV APNs and HLA/CMV APNs carrying αBCMA CAR mRNA was analyzed via flow cytometry. One-way ANOVA with Tukey’s post-test and correction for multiple comparisons; means ± SD, n= 7 biological replicates. ****P<0.0001. **(D)** NSG mice were intravenously inoculated with luciferized human BCMA+ U266 MM tumor cells, injected with enriched human IAV-specific T cells, and dosed with HLA/IAV APNs encapsulated with αBCMA CAR mRNA. Tumor burden was measured via luminescence by IVIS **(E)** and quantified up to 30 days after tumor inoculation **(F)**. Two-way ANOVA with Sidak post-test and correction for multiple comparisons; means ± SEM, n=5 biological replicates. NS = not significant, ***P<0.001, ****P<0.0001.

We further tested whether the APN-transfected *in vivo* CAR T cells would be functional against MM in NSG mice. To test this, we systemically inoculated NSG mice with BCMA+ U266 cancer cells, which were luciferized to allow tracking of tumor growth kinetics by live animal imaging using an IVIS Spectrum CT system. At 6 days after tumor inoculation, we adoptively transferred enriched IAV-specific T cells to tumor-bearing mice, followed by intravenous APN injection at 24 hours after cell transfer (**Fig. 5D**). A total of 5 doses were given to the mice every 5 days. We included αBCMA CAR T cells transduced by lentivirus *ex vivo* (*ex vivo* CAR) for comparison. Notably, we matched the *ex vivo* CAR dose with *in vivo* APN-transfected CAR (∼2e6 CAR+ T cell at 24 hr after APN transfection) and this *ex vivo* CAR dose was lower than curative doses of αBCMA CAR T cells^57^. In the group treated with APNs, only 2 out of 5 mice showed visible bioluminescence signal from U266 at 30 days after tumor inoculation (**Fig. 5E**). Similar to *ex vivo* CAR, APNs carrying αBCMA CAR mRNA resulted in significant tumor regression as compared to the PBS control group for at least 30 days (**Fig. 5F**). Collectively, our data indicated the potential of APNs for redirecting virus-specific T cells *in vivo* for anti-cancer CAR T cell therapy.

### Outlook

The promise of Ag-specific T cell immunotherapy is the potential to tailor T cell responses to specific antigens while reducing off-target effects. This potential can be realized through the unique advantages of mRNA delivery for engineering cell functions. mRNA possess no risk for insertional mutagenesis, does not need to access the nucleus for functionality, and triggers longer lasting expression of therapeutic proteins compared to protein/peptide drugs alone^58^. Combining these qualifications, here we developed APNs that leverage TCR-mediated transfection for Ag-specific immunotherapy by selectively depleting autoreactive T cells and engineering virus-specific T cells with anti-cancer CARs.

Selective depletion of autoreactive Ag-specific T cells to combat autoimmune diseases has been explored using pMHC tetramers conjugated with toxin^59, 60^, lentiviral vectors encoding pro-apoptotic Casp9 transgene^61^, and engineered T cells targeting autoreactive T cells^62^. While promising, the translation potential of a bacterial-derived streptavidin used in pMHC tetramers and lentiviral vectors could be limited by their immunogenicity, whereas broad patient assesses to engineered T cells is hindered by their complex manufacturing process. Alternatively, we demonstrated the development of APNs to selectively deliver pro-apoptotic Casp6 mRNA to induce apoptosis in autoreactive T cells, thereby preventing the onset of hyperglycemia in an aggressive T1D mouse model. Despite the *in vivo* delivery of potent pro-apoptotic Casp6 mRNA afforded by APNs, we did not observe sign of acute toxicity in mice treated with cognate K^d^/NRP-V7 Casp6 APNs. This is consistent with the high transfection specificity of APNs that we reported previously^27^. Our work reported here supports depleting autoreactive T cells by APNs to prevent hyperglycemia in an adoptive cell transfer mouse model of T1D.

*In vivo* production of CAR T cells is emerging as a promising approach to address the manufacturing challenges facing the current FDA-approved CAR T cells, including high costs and long vein-to-vein times, severely restricting patient access associated with the *ex vivo* manufacturing procedures^63^. Examples include the use of viral vectors^14, 15, 16^ or polymeric/lipid nanoparticles^3, 64, 65^ surface-conjugated with pan-T cell antibodies (e.g., αCD3, αCD5) to deliver CAR in the form of DNA or mRNA to circulating T cells. Our results complement these emerging *in vivo* CAR strategies by *in vivo* programming virus-specific T cells, including IAV and CMV, with CAR mRNA. Progress has been made to program virus-specific T cells with CAR transgenes *ex vivo* to take advantage of their memory phenotype for prolonged persistence *in vivo*^29^ and allows vaccination to enhance CAR T cell activity through their native T cell receptors^66^ with a clinical trial (NCT01953900) underway to test the combination of CAR T cells and viral vaccination. The ability of APNs to deliver mRNA to virus-specific T cells *in vivo* enables the re-direction of IAV-specific T cells with functional αBCMA CAR that resulted in tumor regression in NSG mice bearing human MM. Moreover, the ability to expand and transfect virus-specific T cells from a MM patient with CAR mRNA by APNs supports the translational potential of APNs for *in vivo* CAR T cell therapy.

In summary, our results demonstrate the potential of APNs for therapeutic interventions by *in vivo* mRNA delivery to Ag-specific T cells. With the customizable components of APNs, namely pMHC and mRNA, we envision that APNs may be a useful Ag-specific immunotherapy with wide applications in cancer, autoimmune disorders, and infectious diseases.

## Methods

### mRNA synthesis by in vitro reverse transcription

Codon-optimized mRNA for anti-human BCMA CAR and mouse reverse Casp6 for in vivo studies were manufactured by ARNAV Biotech. The mRNA constructs were fully substituted with the modified N1-methyl-pseudouridine. mRNA encoding membrane-anchored nLuc luciferase was a gift from Dr. Philip J. Santangelo (Georgia Tech). mRNA encoding proapoptotic proteins (BIM, BID, mouse granzyme B, mouse reverse caspase 6, mouse dimeric caspase 9, and human reverse caspase 6) for *in vitro* assays were synthesized in house using in vitro transcription (IVT). Briefly, DNA fragments containing the construct sequence were ordered from Integrated DNA Technology and cloned into a plasmid backbone. Plasmid templates were linearized with SpeI-HF (New England Biolabs) overnight at 37^°^C, purified by phenol-chloroform extraction and resuspended in nuclease-free water. IVT was performed using T7 mScript™ Standard mRNA Production System (Cellscript C-MSC11610) following the manufacturer’s protocol with complete substitution of uridine with N1-methyl-pseudouridine. The RNA products were capped in the presence of 2’-O-Methyltransferase to generate Cap-1 structures and the poly(A) tailing reaction was performed for 30 minutes at 37^°^C. The mRNA products were precipitated using lithium chloride (Invitrogen) and resuspended in RNA storage buffer (Invitrogen AM7000). The concentrations of mRNAs were measured using Nanodrop and stored at −80°C. The size of the mRNAs was verified by denaturing RNA gel electrophoresis.

### Caspase mRNA design

Mouse reverse cas6 construct were designed as previously described for human reverse casp6 (ref^67^). Briefly, the large subunit, linker and large unit region of mouse Casp6 was first identified (reference). The small subunit mouse caspase 6 was placed between two Casp6 substrate sequences “MVEID” and “LEHHHHHHVEIDGGSP”. The sequence is followed by the endogenous mouse Casp6 linker extended with GS linkers “GGGGSGGGGSGGGGSGGGGSMTETD” to increase the distance between two subunits. The protein ended with the large subunit of mouse caspase 6. The protein sequences were converted to DNA sequence and codon optimized for murine expression using codon optimization tool (IDT).

### APN preparation and characterization

The procedures for pMHCI expression and APN preparation have been described in detail previously^27^. All lipids were purchased from Avanti Polar Lipids. All the LNP formulations tested were listed in **Table. S1**. Briefly, lipid mixture in ethanol was combined with three volumes of mRNA in acetate buffer (16:1 w/w lipid to mRNA) and injected into micro fluidic mixing device Ignite (Precision Nanosystems) at a total flow rate of 12 ml/min (3:1 flow rate ratio aqueous buffer to ethanol. The resultant LNPs were diluted 40X in PBS and concentrated down using Amicon spin filter (10kDa, Millipore). The total lipid concentration of the concentrated LNPs were measured using Amplex™ Red Cholesterol Assay Kit (Thermo Fisher A12216) and calculated under the assumption that the cholesterol percentage in total lipids remained the same before and after the LNP formation.

To functionalize the synthesized LNPs with pMHC, pMHC was coupled with DSPE-PEG-maleimide and transferred to LNP via post-insertion ^68, 69^. Briefly, a lipid solution of DSPE-PEG (2000)-maleimide was dried under nitrogen and placed in vacuum chamber for 1 h to form a thin film. Lipids were rehydrated in HEPES buffer containing 1 mM EDTA at 20 mg/ml in a 60^°^C water bath for 15 min and sonicated in an ultrasonic bath (Branson) for 5 min. Refolded Cys-terminated pMHCI monomers were reduced with TCEP (1:1 pMHC to TCEP molar ratio) at 37^°^C for 20 min and mixed with the DSPE-PEG (2000)-maleimide solution at 1:5 pMHC:maleimide molar ratio. The conjugation was carried on at R.T. for 5 hours. Lipid-conjugated pMHCI molecules were incubated with the preformed LNPs at 1:3 pMHC:lipid molar ratio at RT for 2 h to incorporate pMHCI onto LNPs. The resultant post-insertion mixture was purified using Sepharose CL-6B gel filtration columns (G-Biosciences, 76361-752).

The sizes of APNs in PBS were measured by dynamic light scattering with Malvern nano-ZS Zetasizer (Malvern). Final lipid concentration was quantified using a phospholipid assay kit (Sigma). The concentration of conjugated pMHCI was determined by BCA assay kit (Sigma). The mRNA encapsulation efficiency was quantified by Quant-iT RiboGreen RNA assay (Life Technology) as previously described ^70^.

### In vitro transfection of NOD8.3 CD8 T cells

NOD8.3 CD8 T cells were isolated from splenoyctes following manufacturer’s protocol (Miltenyi Biotech 130-104-075) and αCD3/αCD28 antibody (BD Biosciences 553057/ 553294) activated for 48 hours. *In vitro* transfection with K^d^/Ctrl (non-cognate) APNs, K^d^/NRP-V7 (cognate) APNs, or LNPs was dosed at 2 µg mRNA/million cells. After 24 hours, transfection readout was measured using flow staining (for VHH and GFP expression) or IVIS (for nLuc expression). To block LDLR, T cells were transfected by APNs in the presence of αLDLR or isotype concentration at 1 µg/ml concentration throughout the 24 hr transfection duration. For the TCR inhibition assay using dasatinib (Sigma Aldrich SML2589-50MG), activated T cells were pre-treated with 50 nM dasatinib for 30 min prior to APN transfection^35^. Dasatinib was also maintained at 50 nM during the 24 hour APN transfection.

### Human IAV-specific T cell expansion

PBMCs from HLA-A2 positive donors were used for in vitro T cell expansion by peptide pulse^71^. PBMCs were cultured in complete CTS OpTmizer medium (CTS OpTmizer T Cell Expansion SFM with CTS supplement A1048501, substituted with L-glutamine, penicillin–streptomycin and 2% human serum, Sigma-Aldrich, H3667) in the presence of the HLA A2.1–restricted influenza matrix peptide (sequence GILGFVFTL, 1 μg/ml), rIL-2 (NIH ICI, 50 IU/ml), rIL-7 (NIH NCI, 25 ng/ml) and rIL-15 (NIH NCI, 25 ng/ml). Flu peptide was only added on the first day of culture, whereas cytokines were supplemented whenever cells were split during the two-week expansion period. At day 14 of cell culture, the frequency of IAV-specific T cells in the total expanded cells were characterized by flow staining and prepared for adoptive cell transfer.

### In vitro transfection and killing assay of human IAV-specific T cells with αBCMA CAR by APNs

To transfect human IAV-specific T cells, APNs were dosed at 2µg mRNA per million cells unless specified otherwise. Treated cells were cultured at 500,000 cells/mL of human T cell media containing 100 units/ml hIL-2. After 24 hours, CAR expression on IAV-specific T cells was determined by flow staining with tetramer and PE-conjugated recombinant BCMA proteins (Acro Biosystem, BCA-HP2H2). *In vitro* killing assays were then performed by mixing APN-transfected BCMA CAR T cells with fLuc transduced U266 MM cells at effector to target (E:T) ratio of 1:1, 2.5:1, and 5:1. The cells were cultured in human T cell media containing 100 units/ml hIL-2. After 24 hour incubation, d-luciferin (Fisher LUCK-2G; 150 µg/ml read concentration) was added to the samples to determine the cytotoxicity of APN-transfected BCMA CAR T cells by IVIS imaging. Maximum cytotoxicity was defined as luminescent signal from wells containing only media, while no cytotoxicity was defined by wells containing only target cells.

### Lentiviral production and transduction of primary human T cells with αBCMA CAR

Lentiviral vectors encoding anti-BCMA CAR construct were either made in house or purchasing from BPS Bioscience (78655). VSV-G pseudotyped lentivirus was produced via transfection of HEK293 T cells (ATCC, CRL-3216) using psPAX2 (Addgene 12260) and pMD2.G (Addgene 12259); viral supernatant was concentrated using PEG-it virus precipitation solution (System Biosciences LV825A-1). For viral transductions of primary human T cells, frozen PBMCs were thawed, incubated at 37°C for 48 h prior to CD3 isolation and activation with human T-Activator Dynabeads (Life Technologies 11131D) at a 3:1 bead:cell ratio for 24 h. To transduce the activated T cells, concentrated lentivirus (MOI: 25) was added to non-tissue culture treated 24-well plates that were coated with retronectin (Takara T100B) according to the manufacturer’s instructions and spun at 1,200 × g for 90 min at room temperature. Subsequently, viral solution was removed and 0.25 ml of human T cells (2.5 × 10^5^ cells per ml) in human T cell media containing 50 units per ml hIL-2 was added to the wells and spun at 1,200 × g for another 60 min at 37 °C. Cells were then incubated on the virus-coated plate for 24 h before expansion, and the Dynabeads were removed at 9 d after T cell activation.

### Animal studies

Male NSG mice and female NOD8.3 (NOD.Cg-Tg(TcraTcrbNY8.3)1Pesa/DvsJ) mice were bred and housed in the Georgia Tech Department of Animal Resources (GT DAR) before use at an age of 8-16 weeks. NOD (NOD/shiltJ) (female, 6-12 weeks old) mice were purchased from Jackson Laboratories. All animal protocols were approved by Georgia Tech Institutional Animal Care and Use Committee (protocols no. A100190, A100191, and A100572). All authors complied with relevant ethical regulations while conducting this study.

### In vivo therapy study using adoptive cell transfer model of type 1 diabetes

The procedures for diabetes induction have been described previously^72^. In brief, splenocytes were isolated from NOD8.3 mice and stimulated with 1 μM NRP-V7 (KYNKANVFL) peptide at 2 × 10^6^ cells/mL for 3 days, washed with PBS, and intravenously injected into host NOD mice (age 5-10 wks) at 15 × 10^6^ cells/mouse. One day after injection of NOD8.3 T cells, mice were treated with the following groups: (1) PBS, (2) K^d^/NRP-V7 APNs loaded with mCasp6 mRNA, (3) K^d^/NRP-V7 APNs loaded with VHH mRNA (mRNA control), (4) K^d^/Ctrl APNs (Kd-pMHC with the non-relevant PR8 peptide TYQRTRALV) loaded with mCasp6 mRNA (non-cognate control), (5) Fc-silent anti-CD3 mAb [145-2C11] (Absolute Antibody, Ab00105-1.4). APNs were dosed at 0.1 mg/kg 1 day and 4 days after injection of NOD8.3 T cells and anti-CD3 mAb was dosed at 2.5 mg/kg daily for 5 days post injection of NOD8.3 T cells (positive treatment control) or 0.1 mg/kg 1 day and 4 day after injections of NOD8.3 T cells. Blood glucose was monitored daily following T cell infusion using blood glucose meters (FreeStyle Freedom Lite, Abbott). Mice were considered diabetic at a blood glucose level >250 mg/dL^73^ and euthanized when blood glucose levels exceeded >400 mg/dL two days in a row. Long-term blood glucose levels were measured 2-3 times weekly until the mice reached 12 weeks of age, which is when spontaneous diabetes may develop due to the NOD background.

### T cell isolation for immunofluorescence staining

The pancreas, pancreatic draining lymph node, spleen, and peripheral blood were isolated from mice at endpoint. The spleen and pancreatic draining lymph node were mechanically disrupted with frosted microscope slides, strained through a 40-μm filter, and the red blood cells (RBCs) were lysed with 1X RBC lysis buffer (BioLegend 420301) for 5 mins on ice and then quenched with 1X PBS. The cells were washed with cold RPMI + 10% fetal bovine serum (FBS).

The pancreas was isolated and digested as described previously^74^. In short, the pancreas was resected and perfused with 2 mL of collagenase XI (0.4 mg/mL) (Sigma C7657) and DNAseI (10 u/mL) (Sigma 10104159001). The pancreas was minced with scissors and incubated at 37 C with a total of 6 mL of collagenase/DNAse I solution for 18 minutes. After gently vortexing for 30 seconds, the solution was quenched with 9 mL of RPMI + 10% FBS, strained through a 40-μm filter, and centrifuged at 1000 x g for 5 mins. The pellet was washed with RPMI + 10% FBS, filtered again through a 40-μm filter, and RBCs were lysed as described above.

Peripheral blood was obtained via cheek bleed or terminally through heart puncture. Up to 500 uL of blood were collected in EDTA blood collection tubes (MiniCollect 450532) and inverted 5 times to prevent clotting. Blood was transferred into 5 mLs of 1X RBC lysis buffer, vortexed for 5 seconds, and incubated at room temperature for 5 minutes. RBC lysis was quenched with 8 mL of cold RPMI + 10% FBS and the cells were strained through a 40-μm filter before plating for cell staining.

### Toxicity analysis

Blood was collected via cardiac puncture and transferred into CAT serum collection tubes (Minicollect 450472). Blood serum was isolated following the manufacturer’s protocol and outsourced to Antech Diagnostics, Inc., for biochemical analysis.

### In vivo transfection of IAV-specific T cells by APNs in NSG mice

8-to 12-week-old male NSG mice were injected with IAV-specific T cells expanded from human PBMC (15e6 PBMC, ∼2e6 IAV-specific T cells per mouse) by intravenous injection. At 24 hour after adoptive cell transfer, mice were intravenously injected with one of the following treatments at 0.5 mg/kg mRNA dose: (1) HLA/IAV APNs loaded with αBCMA CAR mRNA, (2) HLA/IAV APNs loaded with αCD19 CAR mRNA cells (mRNA control), (3) HLA/CMV APNs loaded with αBCMA CAR mRNA cells (non-cognate control), and (4) PBS (carrier control). Splenocytes were harvested at 24 hr later and stained against αCD8 mAb, PE-conjugated recombinant BCMA (Acro Biosystem, BCA-HP2H2), pMHC tetramers (streptavidin, 2 μg/ml) on ice for 30 min. The working concentrations of antibodies were listed in **Table S2**. Epitope pMHC tetramers for staining were obtained from the NIH tetramer core.

### In vivo therapy study using NSG mice with systemic U266 tumor

8-to 12-week-old male NSG mice were irradiated with 200 cGy 1 day prior to intravenous injection of 2×10^6^ U266 human MM cells. U266 cells were transduced to constitutively express luciferase for evaluating tumor burden by quantifying the bioluminescence generated by live U266 cells using IVIS. Fluc activity was measured using an IVIS Spectrum CT (PerkinElmer) 10 min after intravenous injections after intraperitoneal injection of d-luciferin (15 mg/mL, 200 μL per mouse). Animals were randomized based on total body bioluminescence 5 days after tumor cell injection. IAV-specific T cells expanded from human PBMC were injected to the mice (15e6 PBMC, ∼4e6 IAV-specific T cells per mouse) on 6 days after tumor inoculation. Mice treated with *ex vivo* CAR did not receive IAV-specific T cells. At seven days after tumor inoculations, mice were intravenously injected with one of the following treatments: (1) HLA/IAV APNs loaded with αBCMA CAR mRNA, (2) lentivirally-transduced αBCMA CAR T cells (*ex vivo* CAR), and (3) PBS (carrier control). To account for the transient expression of CAR mRNA, A total of 5 doses of APN were injected to mice every 5 days. Animals were euthanized if they exhibited disease model-specific endpoints such as hind-leg paralysis or ruffled.

### Software and statistical analysis

Significant differences between control and treatment groups were determined by various statistical analyses. Student’s t test was used for two groups comparison. One-way analysis of variance (ANOVA) was used for multiple groups comparison. Two-way ANOVA was used when there were subgroups in each group. Data represent means ± SD or SEM in each figure and table as indicated. Statistical analyses were performed using GraphPad Prism 8.0.2 software (GraphPad Software). *P < 0.05, **P < 0.01, ***P < 0.001, and ****P < 0.0001. Flow cytometry data were collected with Cytek Aurora and Cytek Northern Lights, followed by analyzed using FlowJo. In vitro luminescent data were collected with Gen5 2.07 (Biotek). In vivo luminescence data were collected and analyzed with Living Image 4.4.5 (PerkinElmer). Tumor growth curves in vivo were analyzed by two-way ANOVA. Figures were designed in Adobe Illustrator.

## Acknowledgements

The authors thank NIH NCI’s Biological Resources Branch (BRB) Preclinical Biologics Repository for providing cytokines, including recombinant human IL2, IL7, and IL15. The following reagent(s) was obtained through the NIH Tetramer Core Facility: HLA-A*02:01 Influenza A M1 58-66 GILGFVFTL Brilliant Violet 421-Labeled Tetramer and HLA-A*02:01 CMV pp65 495-503 NLVPMVATV Brilliant Violet 421-Labeled Tetramer. This work was supported in part by NIH grants R21CA260247; NSF grant ECCS-1542174; The International Myeloma Society (IMS) and Paula and Rodger Riney Foundation Translational Research Award; and the Donaldson Charitable Trust Research Synergy Fund (G.A.K.). F.Y.S. acknowledges support from the National Institute of Health Pathway to Independence Award (K99CA276890) and the Postdoctoral Fellowship jointly provided by the Department of Biomedical Engineering at Georgia Tech and the College of Engineering at Peking University, China. J.C.S., M. Y. W., and A.D.S.T. were supported by the NSF Graduate Research Fellowships Program (Grant No. DGE-2039655). A.D.S.T. was also supported by the NIH Cell and Tissue Engineering (CTEng) Training Program (5T32GM8433-30). This content is solely the responsibility of the authors and does not necessarily represent the official views of the National Institutes of Health.

## Author Information

These authors contributed equally: F.Y.S., J.C.S., C.S.C.

## Author Contributions

F.Y.S., J.C.S., C.S.C. and G.A.K. conceived the idea. F.Y.S., J.C.S., C.S.C., M.Y.W. and G.A.K. designed experiments and interpreted results. F.Y.S., J.C.S., C.S.C., M.Y.W., X.Y., A.S.T., M.S., R.H., N.S., C.H.N., A.G., and J.M. synthesized materials and carried out the experiments. F.Y.S., J.C.S., C.S.C. and G.A.K. wrote the manuscript.

## Reference

1. Breda L, Papp TE, Triebwasser MP, Yadegari A, Fedorky MT, Tanaka N, et al. In vivo hematopoietic stem cell modification by mRNA delivery. Science 2023, 381(6656): 436–443.

2. Hamilton JR, Chen E, Perez BS, Sandoval Espinoza CR, Kang MH, Trinidad M, et al. In vivo human T cell engineering with enveloped delivery vehicles. Nature Biotechnology 2024.

3. Parayath NN, Stephan SB, Koehne AL, Nelson PS, Stephan MT. In vitro-transcribed antigen receptor mRNA nanocarriers for transient expression in circulating T cells in vivo. Nature communications 2020, 11(1): 6080.

4. Rurik JG, Tombácz I, Yadegari A, Méndez Fernández PO, Shewale SV, Li L, et al. CAR T cells produced in vivo to treat cardiac injury. Science 2022, 375(6576): 91–96.

5. Mullard A. FDA approves first BCMA-targeted CAR-T cell therapy. Nature reviews Drug discovery 2021, 20(5): 332.

6. Mullard A. FDA approves first TCR-engineered T cell therapy, for rare soft-tissue cancer. Nature reviews Drug discovery 2024.

7. Wang S, Du Y, Zhang B, Meng G, Liu Z, Liew SY, et al. Transplantation of chemically induced pluripotent stem-cell-derived islets under abdominal anterior rectus sheath in a type 1 diabetes patient. Cell 2024.

8. Siebart JC, Chan CS, Yao X, Su F-Y, Kwong GA. In vivo gene delivery to immune cells. Current opinion in biotechnology 2024, 88: 103169.

9. Mullard A. In vivo CAR T cells move into clinical trials. Nature reviews Drug discovery 2024.

10. Xu EJK, Smith BE, Conce Alberto WD, Walsh MJ, Lim B, Hoffman MT, et al. Peptide-MHC-targeted retroviruses enable in *vivo* expansion and gene delivery to tumor-specific T cells. bioRxiv 2024: 2024.2009.2018.613594.

11. Nahmad AD, Lazzarotto CR, Zelikson N, Kustin T, Tenuta M, Huang D, et al. In vivo engineered B cells secrete high titers of broadly neutralizing anti-HIV antibodies in mice. Nature Biotechnology 2022, 40(8): 1241–1249.

12. Kerzel T, Giacca G, Beretta S, Bresesti C, Notaro M, Scotti GM, et al. In *vivo* macrophage engineering reshapes the tumor microenvironment leading to eradication of liver metastases. Cancer Cell 2023, 41(11): 1892–1910.e1810.

13. Ascic E, Åkerström F, Sreekumar Nair M, Rosa A, Kurochkin I, Zimmermannova O, et al. In vivo dendritic cell reprogramming for cancer immunotherapy. Science, 0(0): eadn9083.

14. Nawaz W, Huang B, Xu S, Li Y, Zhu L, Wu Z, et al. AAV-Mediated In Vivo CAR Gene Therapy for Targeting Human T Cell Leukemia. bioRxiv 2021: 2021.2002.2015.431201.

15. Venditti CP. Safety questions for AAV gene therapy. Nature Biotechnology 2021, 39(1): 24–26.

16. Shirley JL, de Jong YP, Terhorst C, Herzog RW. Immune Responses to Viral Gene Therapy Vectors. Molecular Therapy 2020, 28(3): 709–722.

17. Lee DY, Amirthalingam S, Lee C, Rajendran AK, Ahn YH, Hwang NS. Strategies for targeted gene delivery using lipid nanoparticles and cell-derived nanovesicles. Nanoscale Adv 2023, 5(15): 3834–3856.

18. Billingsley MM, Gong N, Mukalel AJ, Thatte AS, El-Mayta R, Patel SK, et al. In Vivo mRNA CAR T Cell Engineering via Targeted Ionizable Lipid Nanoparticles with Extrahepatic Tropism. Small 2024, 20(11): e2304378.

19. Porciello N, Franzese O, D’Ambrosio L, Palermo B, Nisticò P. T-cell repertoire diversity: friend or foe for protective antitumor response? Journal of Experimental & Clinical Cancer Research 2022, 41(1): 356.

20. Quandt Z, Young A, Perdigoto AL, Herold KC, Anderson MS. Autoimmune Endocrinopathies: An Emerging Complication of Immune Checkpoint Inhibitors. Annual review of medicine 2021, 72: 313–330.

21. Gaston RS, Deierhoi MH, Patterson T, Prasthofer E, Julian BA, Barber WH, et al. OKT3 first-dose reaction: Association with T cell subsets and cytokine release. Kidney International 1991, 39(1): 141–148.

22. Keymeulen B, Candon S, Fafi-Kremer S, Ziegler A, Leruez-Ville M, Mathieu C, et al. Transient Epstein-Barr virus reactivation in CD3 monoclonal antibody-treated patients. Blood 2010, 115(6): 1145–1155.

23. Guo X-zJ, Elledge SJ. V-CARMA: A tool for the detection and modification of antigen-specific T cells. Proceedings of the National Academy of Sciences 2022, 119(4): e2116277119.

24. Dobson CS, Reich AN, Gaglione S, Smith BE, Kim EJ, Dong J, et al. Antigen identification and high-throughput interaction mapping by reprogramming viral entry. Nat Methods 2022, 19(4): 449–460.

25. Dahotre SN, Romanov AM, Su F-Y, Kwong GA. Synthetic Antigen-Presenting Cells for Adoptive T Cell Therapy. Advanced Therapeutics 2021, 4(8): 2100034.

26. Quayle SN, Girgis N, Thapa DR, Merazga Z, Kemp MM, Histed A, et al. CUE-101, a Novel E7-pHLA-IL2-Fc Fusion Protein, Enhances Tumor Antigen-Specific T-Cell Activation for the Treatment of HPV16-Driven Malignancies. Clinical cancer research : an official journal of the American Association for Cancer Research 2020, 26(8): 1953–1964.

27. Su F-Y, Zhao QH, Dahotre SN, Gamboa L, Bawage SS, Silva Trenkle AD, et al. In vivo mRNA delivery to virus-specific T cells by light-induced ligand exchange of MHC class I antigen-presenting nanoparticles. Science Advances 2022, 8(8): eabm7950.

28. Yang K, Zhang Y, Ding J, Li Z, Zhang H, Zou F. Autoimmune CD8+ T cells in type 1 diabetes: from single-cell RNA sequencing to T-cell receptor redirection. Front Endocrinol (Lausanne) 2024, 15: 1377322.

29. Rosato PC, Wijeyesinghe S, Stolley JM, Nelson CE, Davis RL, Manlove LS, et al. Virus-specific memory T cells populate tumors and can be repurposed for tumor immunotherapy. Nature communications 2019, 10(1): 567.

30. Wang Y, Miao L, Satterlee A, Huang L. Delivery of oligonucleotides with lipid nanoparticles. Advanced drug delivery reviews 2015, 87: 68–80.

31. Lieberman SM, Evans AM, Han B, Takaki T, Vinnitskaya Y, Caldwell JA, et al. Identification of the beta cell antigen targeted by a prevalent population of pathogenic CD8+ T cells in autoimmune diabetes. Proc Natl Acad Sci U S A 2003, 100(14): 8384–8388.

32. Kim M, Jeong M, Hur S, Cho Y, Park J, Jung H, et al. Engineered ionizable lipid nanoparticles for targeted delivery of RNA therapeutics into different types of cells in the liver. Science Advances 2021, 7(9): eabf4398.

33. Akinc A, Querbes W, De S, Qin J, Frank-Kamenetsky M, Jayaprakash KN, et al. Targeted Delivery of RNAi Therapeutics With Endogenous and Exogenous Ligand-Based Mechanisms. Molecular Therapy 2010, 18(7): 1357–1364.

34. Goyette J, Nieves DJ, Ma Y, Gaus K. How does T cell receptor clustering impact on signal transduction? Journal of Cell Science 2019, 132(4).

35. Lissina A, Ladell K, Skowera A, Clement M, Edwards E, Seggewiss R, et al. Protein kinase inhibitors substantially improve the physical detection of T-cells with peptide-MHC tetramers. Journal of immunological methods 2009, 340(1): 11–24.

36. Herold KC, Delong T, Perdigoto AL, Biru N, Brusko TM, Walker LSK. The immunology of type 1 diabetes. Nature Reviews Immunology 2024, 24(6): 435–451.

37. Long SA, Thorpe J, DeBerg HA, Gersuk V, Eddy JA, Harris KM, et al. Partial exhaustion of CD8 T cells and clinical response to teplizumab in new-onset type 1 diabetes. Science Immunology 2016, 1(5): eaai7793.

38. Herold KC, Bundy BN, Long SA, Bluestone JA, DiMeglio LA, Dufort MJ, et al. An Anti-CD3 Antibody, Teplizumab, in Relatives at Risk for Type 1 Diabetes. The New England journal of medicine 2019, 381(7): 603–613.

39. Kern HB, Srinivasan S, Convertine AJ, Hockenbery D, Press OW, Stayton PS. Enzyme-Cleavable Polymeric Micelles for the Intracellular Delivery of Proapoptotic Peptides. Molecular pharmaceutics 2017, 14(5): 1450–1459.

40. Chao Y, Shiozaki EN, Srinivasula SM, Rigotti DJ, Fairman R, Shi Y. Engineering a dimeric caspase-9: a re-evaluation of the induced proximity model for caspase activation. PLoS biology 2005, 3(6): e183.

41. Srinivasula SM, Ahmad M, MacFarlane M, Luo Z, Huang Z, Fernandes-Alnemri T, et al. Generation of constitutively active recombinant caspases-3 and-6 by rearrangement of their subunits. Journal of Biological Chemistry 1998, 273(17): 10107–10111.

42. Chatenoud L, Thervet E, Primo J, Bach JF. Anti-CD3 antibody induces long-term remission of overt autoimmunity in nonobese diabetic mice. Proc Natl Acad Sci U S A 1994, 91(1): 123–127.

43. Sims EK, Bundy BN, Stier K, Serti E, Lim N, Long SA, et al. Teplizumab improves and stabilizes beta cell function in antibody-positive high-risk individuals. Sci Transl Med 2021, 13(583).

44. Mikkilineni L, Kochenderfer JN. CAR T cell therapies for patients with multiple myeloma. Nat Rev Clin Oncol 2021, 18(2): 71–84.

45. Raje N, Berdeja J, Lin Y, Siegel D, Jagannath S, Madduri D, et al. Anti-BCMA CAR T-Cell Therapy bb2121 in Relapsed or Refractory Multiple Myeloma. The New England journal of medicine 2019, 380(18): 1726–1737.

46. Slaney CY, von Scheidt B, Davenport AJ, Beavis PA, Westwood JA, Mardiana S, et al. Dual-specific Chimeric Antigen Receptor T Cells and an Indirect Vaccine Eradicate a Variety of Large Solid Tumors in an Immunocompetent, Self-antigen Setting. Clinical cancer research : an official journal of the American Association for Cancer Research 2017, 23(10): 2478–2490.

47. Tanaka M, Tashiro H, Omer B, Lapteva N, Ando J, Ngo M, et al. Vaccination Targeting Native Receptors to Enhance the Function and Proliferation of Chimeric Antigen Receptor (CAR)-Modified T Cells. Clin Cancer Res 2017, 23(14): 3499–3509.

48. Wang X, Wong CW, Urak R, Mardiros A, Budde LE, Chang WC, et al. CMVpp65 Vaccine Enhances the Antitumor Efficacy of Adoptively Transferred CD19-Redirected CMV-Specific T Cells. Clin Cancer Res 2015, 21(13): 2993–3002.

49. Su F-Y, Zhao Q, Dahotre SN, Gamboa L, Bawage SS, Silva Trenkle AD, et al. *In vivo* mRNA delivery to virus-specific T cells by light-induced ligand exchange of MHC class I antigen-presenting nanoparticles. bioRxiv 2021: 2021.2010.2014.464373.

50. Simon S, Riddell SR. Dual Targeting with CAR T Cells to Limit Antigen Escape in Multiple Myeloma. Blood Cancer Discov 2020, 1(2): 130–133.

51. Zhou D, Sun Q, Xia J, Gu W, Qian J, Zhuang W, et al. Anti-BCMA/GPRC5D bispecific CAR T cells in patients with relapsed or refractory multiple myeloma: a single-arm, single-centre, phase 1 trial. The Lancet Haematology.

52. Fernández de Larrea C, Staehr M, Lopez AV, Ng KY, Chen Y, Godfrey WD, et al. Defining an Optimal Dual-Targeted CAR T-cell Therapy Approach Simultaneously Targeting BCMA and GPRC5D to Prevent BCMA Escape-Driven Relapse in Multiple Myeloma. Blood Cancer Discov 2020, 1(2): 146–154.

53. Zhang M, Wei G, Zhou L, Zhou J, Chen S, Zhang W, et al. GPRC5D CAR T cells (OriCAR-017) in patients with relapsed or refractory multiple myeloma (POLARIS): a first-in-human, single-centre, single-arm, phase 1 trial. The Lancet Haematology 2023, 10(2): e107–e116.

54. Mailankody S, Devlin SM, Landa J, Nath K, Diamonte C, Carstens EJ, et al. GPRC5D-Targeted CAR T Cells for Myeloma. The New England journal of medicine 2022, 387(13): 1196–1206.

55. Smith EL, Harrington K, Staehr M, Masakayan R, Jones J, Long TJ, et al. GPRC5D is a target for the immunotherapy of multiple myeloma with rationally designed CAR T cells. Sci Transl Med 2019, 11(485).

56. Liu RJBLS. Chimeric antigen receptors targeting G-protein coupled receptor and uses thereof. 2017.

57. Duan D, Wang K, Wei C, Feng D, Liu Y, He Q, et al. The BCMA-Targeted Fourth-Generation CAR-T Cells Secreting IL-7 and CCL19 for Therapy of Refractory/Recurrent Multiple Myeloma. Front Immunol 2021, 12: 609421.

58. Qin S, Tang X, Chen Y, Chen K, Fan N, Xiao W, et al. mRNA-based therapeutics: powerful and versatile tools to combat diseases. Signal Transduction and Targeted Therapy 2022, 7(1): 166.

59. Hess PR, Barnes C, Woolard MD, Johnson MD, Cullen JM, Collins EJ, et al. Selective deletion of antigen-specific CD8+ T cells by MHC class I tetramers coupled to the type I ribosome-inactivating protein saporin. Blood 2007, 109(8): 3300–3307.

60. Goldberg SD, Felix N, McCauley M, Eberwine R, Casta L, Haskell K, et al. A Strategy for Selective Deletion of Autoimmunity-Related T Cells by pMHC-Targeted Delivery. Pharmaceutics 2021, 13(10).

61. Guo XJ, Elledge SJ. V-CARMA: A tool for the detection and modification of antigen-specific T cells. Proc Natl Acad Sci U S A 2022, 119(4).

62. Fishman S, Lewis MD, Siew LK, De Leenheer E, Kakabadse D, Davies J, et al. Adoptive Transfer of mRNA-Transfected T Cells Redirected against Diabetogenic CD8 T Cells Can Prevent Diabetes. Molecular Therapy 2017, 25(2): 456–464.

63. Ayala Ceja M, Khericha M, Harris CM, Puig-Saus C, Chen YY. CAR-T cell manufacturing: Major process parameters and next-generation strategies. The Journal of experimental medicine 2024, 221(2).

64. Rurik JG, Tombácz I, Yadegari A, Fernández POM, Shewale SV, Li L, et al. CAR T cells produced in vivo to treat cardiac injury. Science 2022, 375(6576): 91–96.

65. 61. Effects of Chimeric Antigen Receptor (CAR) Expression on Regulatory T Cells. Molecular Therapy 2009, 17: S25.

66. Slaney CY, von Scheidt B, Davenport AJ, Beavis PA, Westwood JA, Mardiana S, et al. Dual-specific Chimeric Antigen Receptor T Cells and an Indirect Vaccine Eradicate a Variety of Large Solid Tumors in an Immunocompetent, Self-antigen Setting. Clinical Cancer Research 2017, 23(10): 2478–2490.

67. Srinivasula SM, Ahmad M, MacFarlane M, Luo Z, Huang Z, Fernandes-Alnemri T, et al. Generation of Constitutively Active Recombinant Caspases-3 and -6 by Rearrangement of Their Subunits*. Journal of Biological Chemistry 1998, 273(17): 10107–10111.

68. Ishida T, Iden DL, Allen TM. A combinatorial approach to producing sterically stabilized (Stealth) immunoliposomal drugs. FEBS Letters 1999, 460(1): 129–133.

69. Lainé AL, Gravier J, Henry M, Sancey L, Béjaud J, Pancani E, et al. Conventional versus stealth lipid nanoparticles: Formulation and in vivo fate prediction through FRET monitoring. Journal of Controlled Release 2014, 188: 1–8.

70. Kedmi R, Veiga N, Ramishetti S, Goldsmith M, Rosenblum D, Dammes N, et al. A modular platform for targeted RNAi therapeutics. Nature Nanotechnology 2018, 13(3): 214–219.

71. Eberhardt CS, Kissick HT, Patel MR, Cardenas MA, Prokhnevska N, Obeng RC, et al. Functional HPV-specific PD-1+ stem-like CD8 T cells in head and neck cancer. Nature 2021, 597(7875): 279–284.

72. Gammon JM, Carey ST, Saxena V, Eppler HB, Tsai SJ, Paluskievicz C, et al. Engineering the lymph node environment promotes antigen-specific efficacy in type 1 diabetes and islet transplantation. Nature communications 2023, 14(1): 681.

73. Mathews CE, Xue S, Posgai A, Lightfoot YL, Li X, Lin A, et al. Acute Versus Progressive Onset of Diabetes in NOD Mice: Potential Implications for Therapeutic Interventions in Type 1 Diabetes. Diabetes 2015, 64(11): 3885–3890.

74. Gearty SV, Dündar F, Zumbo P, Espinosa-Carrasco G, Shakiba M, Sanchez-Rivera FJ, et al. An autoimmune stem-like CD8 T cell population drives type 1 diabetes. Nature 2022, 602(7895): 156–161.

