## Supplemental Figure 1 for "Antigen-specific T cell immunotherapy by in vivo mRNA delivery"

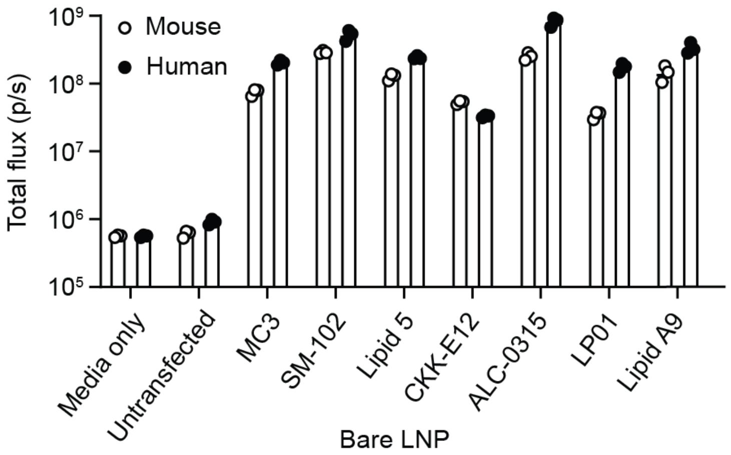


**Fig. S1. Optimization of lipid nanoparticle (LNP) formulation for primary mouse and human T cell transfection.** IVIS quantification of bioluminescence intensity of human and mouse CD8 T cells transfected with various LNP formulations encapsulated with membrane-bound nano-Luciferase mRNA for 24 hours.
