## Supplemental Figure 2 for "Antigen-specific T cell immunotherapy by in vivo mRNA delivery"

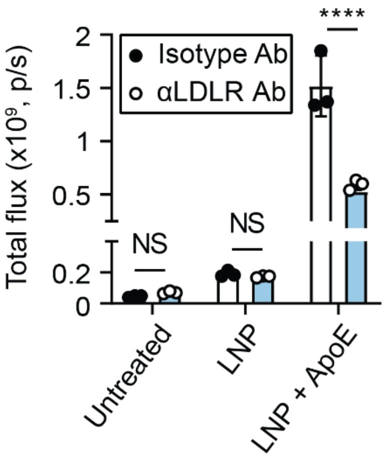


**Fig. S2. T cell transfection by lipid nanoparticle (LNP) is low density lipoprotein receptor (LDLR)-dependent.** Activated NOD8.3 CD8 T cells were transfected with LNPs loaded with nLuc in the presence of either a LDLR blocking antibody (αLDLR Ab) or an isotype antibody control (isotype Ab). Transfection was measured by IVIS 24 hours post transfection.
