## Supplemental Figure 3 for "Antigen-specific T cell immunotherapy by in vivo mRNA delivery"

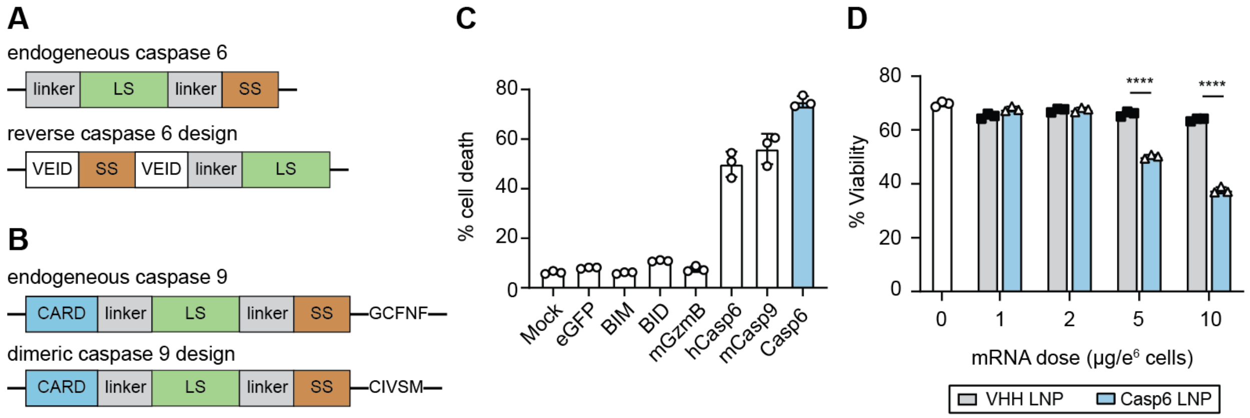


**Fig. S3. Design and characterization of proapoptotic mRNA for depletion of autoreactive T cells. (A)** Schematic detailing endogenous caspase 6 and reverse caspase 6 design. Endogenous Casp6; linker, large subunit (LS), linker, small subunit (SS). Reverse Casp6; VEID amino acid sequence, small subunit (SS), VEID amino acid sequence, linker large subunit (LS). **(B)** Schematic detailing endogenous caspase 9 and dimeric caspase 9 design. Endogenous Casp9; caspase recruitment domain (CARD), linker, large subunit (LS), linker, small subunit (SS), GCFNF amino acid sequence. Dimeric Casp9; caspase recruitment domain (CARD), linker, large subunit (LS), linker, small subunit (SS), CIVSM amino acid sequence. **(C)** Quantification of mouse CD8 T cell death 24 hours after electroporation with mRNA encoding for the specified proapoptotic protein. B-cell Lymphoma 2-like protein 11 (BIM); BH3 interacting-domain death agonist (BID); mouse granzyme B (mGzmB); human caspase 6 (hCasp6); mouse dimeric caspase 9 (mCasp9); mouse reverse caspase 6 (mCasp6). **(D)** Quantification of cell viability after *in vitro* transfection of mouse CD8 T cells with LNPs encapsulated with VHH or mouse Casp6 mRNA. Two-way ANOVA with Sidak post-test and correction for multiple comparisons; means ± SD, n=3 independent wells ****P<0.000.
