## Supplemental Figure 4 for "Antigen-specific T cell immunotherapy by in vivo mRNA delivery"

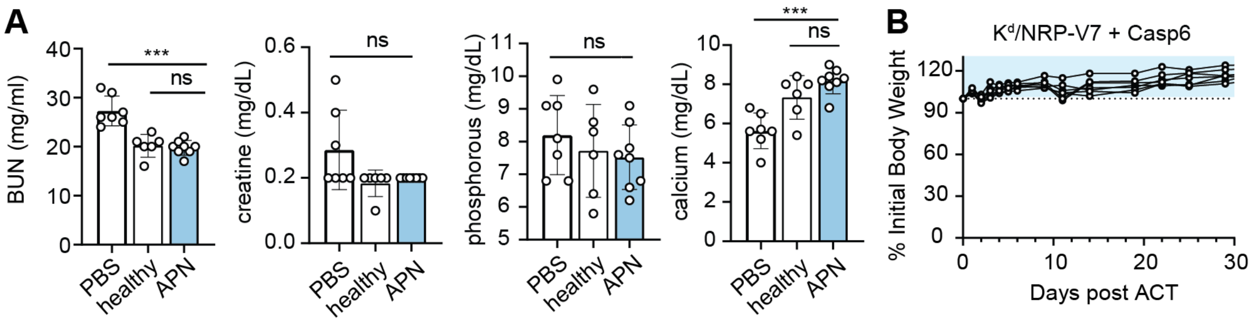


**Fig. S4. Biocompatibility of APNs in murine ACT model of T1D. (A)** Quantification of serum blood urea nitrogen (BUN), creatinine, phosphorous, and calcium after treatment with $K^d$/NRP-V7 Casp6 APNs to deplete NOD8.3 T cells in murine ACT model of T1D. One-way ANOVA with Tukey's post-test and correction for multiple comparisons; means ± SD, n=7-8 biological replicates. ns = not significant, ***P<0.001. **(B)** Quantification of % body weight change after treatment with $K^d$/NRP-V7 Casp6 APNs. Dashed line represents initial body weight and light-blue shaded region shows healthy body weight range.
