## Supplemental Figure 5 for "Antigen-specific T cell immunotherapy by in vivo mRNA delivery"

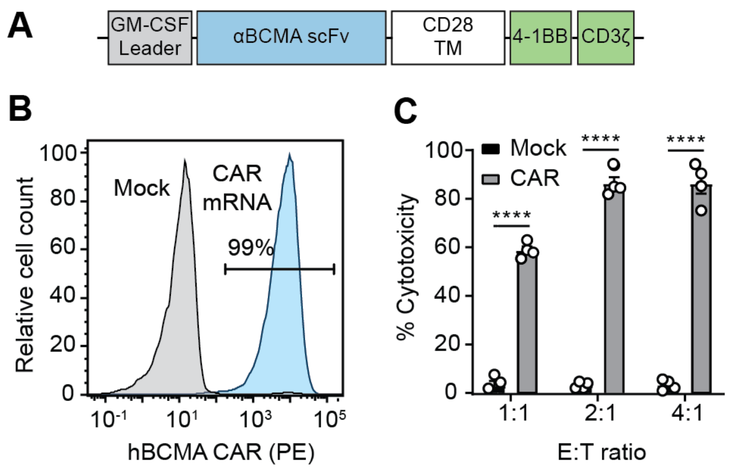


**Fig. S5. Design and characterization of αBCMA CAR mRNA. (A)** Schematic detailing the design of αBCMA CAR mRNA. Granulocyte-macrophage colony-stimulating factor (GM-CSF) leader sequence, αBCMA CAR single-chain variable fragment (αBCMA CAR scFv), CD28 transmembrane domain (CD28 TM), and 4-1BB/CD3ζ. **(B)** Flow cytometry analysis of CAR expression after electroporation with human αBCMA CAR mRNA. **(C)** *In vitro* quantification of % cytotoxicity of electroporated human αBCMA CAR T cells coincubated with luciferized BCMA+ MM1R tumor cells. Two-way ANOVA with Sidak post-test and correction for multiple comparisons; means ± SD, n=4 independent wells. ****P<0.0001.
