## Supplemental Figure 6 for "Antigen-specific T cell immunotherapy by in vivo mRNA delivery"

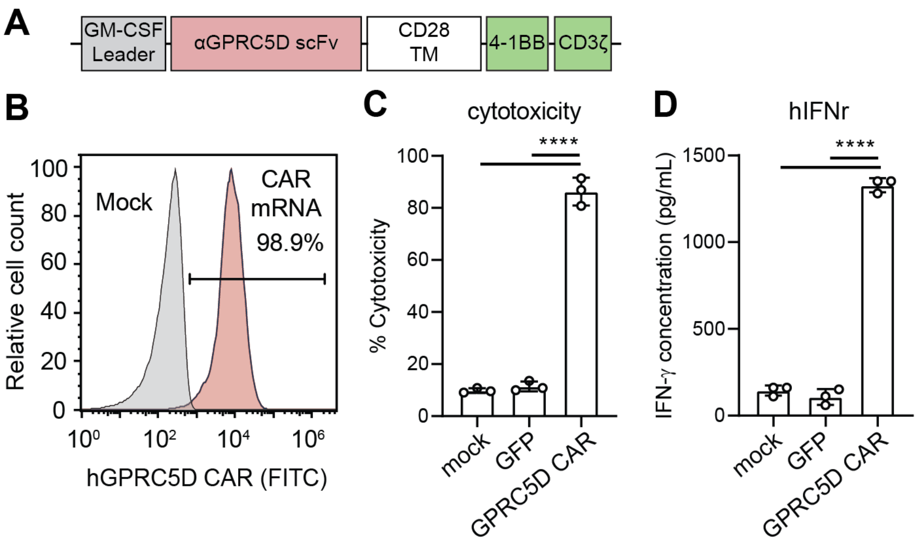


**Fig. S6. Design and characterization of αGPRC5D CAR mRNA. (A)** Schematic detailing design of αGPRC5D CAR mRNA. Granulocyte-macrophage colony-stimulating factor (GM-CSF) leader sequence, αGPRC5D CAR single-chain variable fragment (αBCMA CAR scFv), CD28 transmembrane domain (CD28 TM), and 4-1BB/CD3ζ. **(B)** Flow cytometry analysis of CAR expression after electroporation with human αGPRC5D CAR mRNA. **(C)** *In vitro* quantification of % cytotoxicity of electroporated human αGPRC5D CAR T cells coincubated with luciferized GPRC5D+ MM1R tumor cells. **(D)** Quantification of human IFN-Ƴ detected in the serum of cocultures via ELISA. One-way ANOVA with Tukey's post-test and correction for multiple comparisons; means ± SD, n=3 independent wells. ****P<0.0001.
