## Supplemental Figure 7 for "Antigen-specific T cell immunotherapy by in vivo mRNA delivery"

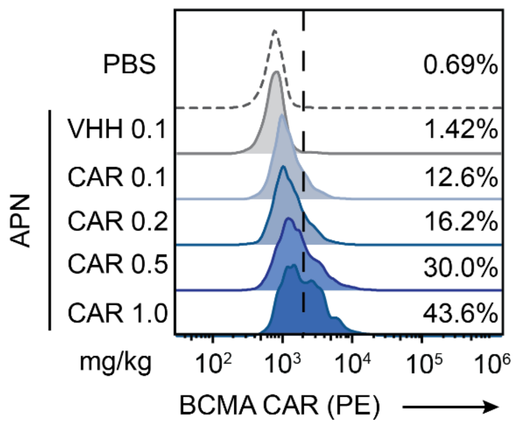


**Fig. S7. APN transfect human IAV-specific T cells with αBCMA CAR mRNA *in vivo* in a dose-dependent manner.** Enriched human IAV-specific T cells were infused in NSG mice and HLA/IAV APNs encapsulated with αBCMA CAR mRNA were intravenously infused 24 hours after. Representative flow plots analyzing *in vivo* transfection efficiency of HLA/IAV APNs at increasing doses.
