## Supplemental Table 1 for "Antigen-specific T cell immunotherapy by in vivo mRNA delivery"

**Table S1. Lipid components of LNP formulations tested in Fig. S1.**

| LNP formulation | ionizable lipid %mol | | cholesterol %mol | phospholipid %mol | | PEG-Lipid %mol | |
|---|---|---|---|---|---|---|---|
| **MC3** | MC3 | 50 | 38.5 | DSPC | 10 | DMG-PEG | 1.5 |
| **SM-102** | SM-102 | 50 | 38.5 | DSPC | 10 | DMG-PEG | 1.5 |
| **Lipid 5** | Lipid 5 | 50 | 38.5 | DSPC | 10 | DMG-PEG | 1.5 |
| **CKK-E12** | CKK-E12 | 35 | 46.5 | DOPE | 16 | PEG14-2000 | 2.5 |
| **ALC-0315** | ALC-0315 | 46.3 | 42.7 | DSPC | 9.4 | ALC-0159 | 1.6 |
| **LP01** | LP01 | 45 | 44 | DSPC | 9 | DMG-PEG | 2 |
| **Lipid A9** | Lipid A9 | 50 | 38.5 | DSPC | 10 | PEG14-2000 | 1.5 |
