## Supplemental Table 2 for "Antigen-specific T cell immunotherapy by in vivo mRNA delivery"

### Table S2. List of staining reagents for flowcytometry analysis

| REAGENT | | | | | |
|---|---|---|---|---|---|
| **Antibodies** | | | | | |
| **Target** | **Fluorochrome** | **Clone** | **Source** | **Identifier** | **Dilution factor** |
| Camelid VHH | iFluor488 | 96A3F5 | Genscript | A01862 | 1:100 |
| Mouse CD8a | APC/Cy7 | 53-6.7 | Biolegend | 100714 | 1:100 |
| Mouse CD3 | APC | 17A2 | Biolegend | 100236 | 1:100 |
| Human CD8a | APC | HIT8a | Biolegend | 300912 | 1:100 |
| Human CD8a | FITC | HIT8a | Biolegend | 300906 | 1:100 |
| Human CD3 | FITC | HIT3a | Biolegend | 300306 | 1:100 |
| Strep II tag | FITC | 5A9F9 | Genscript | A01736-100 | 1:50 |
| **Other reagents** | | | | | |
| $K^d$/NRP-V7 tetramer | PE | | Made in house with PE-conjugated streptavidin (Thermo Fisher S866) | | 2 µg streptavidin/sample |
| HLA-A2/IAV-GIL tetramer | BV421 | | Tetramer core | | 1:50 |
| HLA-A2/CMV-NLV tetramer | BV421 | | Tetramer core | | 1:50 |
| Recombinant BCMA | PE | | Acro Biosystem | BCA-HP2H2 | 1:20 |
| LIVE/DEAD™ Fixable Aqua Dead Cell Stain Kit | | | Invitrogen | L34957 | 1:100 |
| LIVE/DEAD™ Fixable Near IR Dead Cell Stain Kit | | | Invitrogen | L10119 | 1:100 |
